# Sexual size dimorphism increases with body size at the intraspecific level in *Drosophila melanogaster*

**DOI:** 10.64898/2026.01.25.701610

**Authors:** Yang Zhang, Quanquan Jin, Xinqiang Xi

## Abstract

Sexual size dimorphism (SSD), the difference in body size between males and females, typically conforms to Rensch’s rule across species: SSD increases with body size when males are larger but decreases when females are larger. Although this macroevolutionary pattern has been extensively documented, intraspecific analyses remain rare, yet they are essential for understanding the proximate mechanisms underlying the origin and maintenance of sexual dimorphism. In particular, it remains unclear whether within-species variation in SSD is driven primarily by sex-specific differences in growth rate or in development time. Here, we addressed this question by examining SSD scaling in inbred lines of *Drosophila melanogaster* from the well-established *Drosophila melanogaster* Genetic Reference Panel (DGRP) reared under two thermal environments (25 °C and 28 °C). Females were consistently larger than males, resulting in pronounced female-biased SSD across different lines of this model insect. Moreover, SSD increased with overall body size, representing a reversal of Rensch’s rule at the intraspecific level. This scaling pattern was largely explained by higher female growth rates rather than sexual differences in development time. Elevated temperature reduced SSD by decreasing female growth rate while slightly enhancing that of males. Together, our results demonstrate that Rensch’s rule does not universally apply at intraspecific level and underscore the critical role of growth rate and environmental sensitivity in shaping SSD at the intraspecific level.

## INTRODUCTION

Sexual size dimorphism (SSD), differences in body size between males and females, is widespread across the animal kingdom (Abouheif & Fairbairn, 1997; Fairbairn, 1997). A central macroevolutionary pattern describing SSD is Rensch’s rule (Rensch, 1950, 1960), which predicts that SSD increases with mean body size when males are larger, but decreases with body size when females are larger. This rule has been supported in many vertebrate taxa (e.g., Colwell, 2000; Cox et al., 2003; Blanckenhorn et al., 2007), yet notable exceptions occur, including reversed patterns in some invertebrates such as beetles (Blanckenhorn et al., 2007). These deviations imply that the evolution of SSD cannot be fully understood from interspecific comparisons alone.

Importantly, macroevolutionary correlations often have limited mechanistic explanatory power because evolutionary change ultimately reflects selection acting on heritable variation within species (Li et al., 2018). Therefore, a deeper understanding of SSD requires evaluating whether patterns observed among species are mirrored within species (Fairbairn et al., 2007). Intraspecific tests can provide a critical bridge between macroevolutionary patterns and the microevolutionary processes that generate and maintain variation (Stillwell & Fox, 2007, 2009; Tsuboi et al., 2024). Genetic diversity and environmental heterogeneity can produce sex-specific differences in life-history traits, yielding divergent SSD-body size relationships across genotypes and environments (Shine, 1990; Badyaev, 2002). As a result, reliance on species-level mean SSD may oversimplify interspecific patterns and obscure the intraspecific level dynamics underlying dimorphism.

Several hypotheses have been proposed to explain how SSD relates to body size. The sexual selection hypothesis posits that male-biased SSD evolves when larger males gain reproductive advantages through mate competition (Andersson & Iwasa, 1996; Székely et al., 2004). The fecundity selection hypothesis predicts female-biased SSD when larger females achieve higher reproductive output, potentially producing inverse Rensch patterns (Stillwell & Davidowitz, 2010; Barneche et al., 2018). Finally, the sex-specific phenotypic plasticity hypothesis suggests that males and females respond differently to environmental variation (e.g., temperature, resource availability), thereby shifting SSD allometry in either direction (Fairbairn, 2005; Stillwell & Davidowitz, 2010). These alternative predictions can be summarized graphically as a set of potential SSD-body size allometries (Figure 1), highlighting how Rensch versus inverse-Rensch patterns may arise at different evolutionary scales. Distinguishing among these hypotheses benefits from intraspecific approaches that directly link sex-specific growth and environmental responses to SSD.

**Figure 1.**
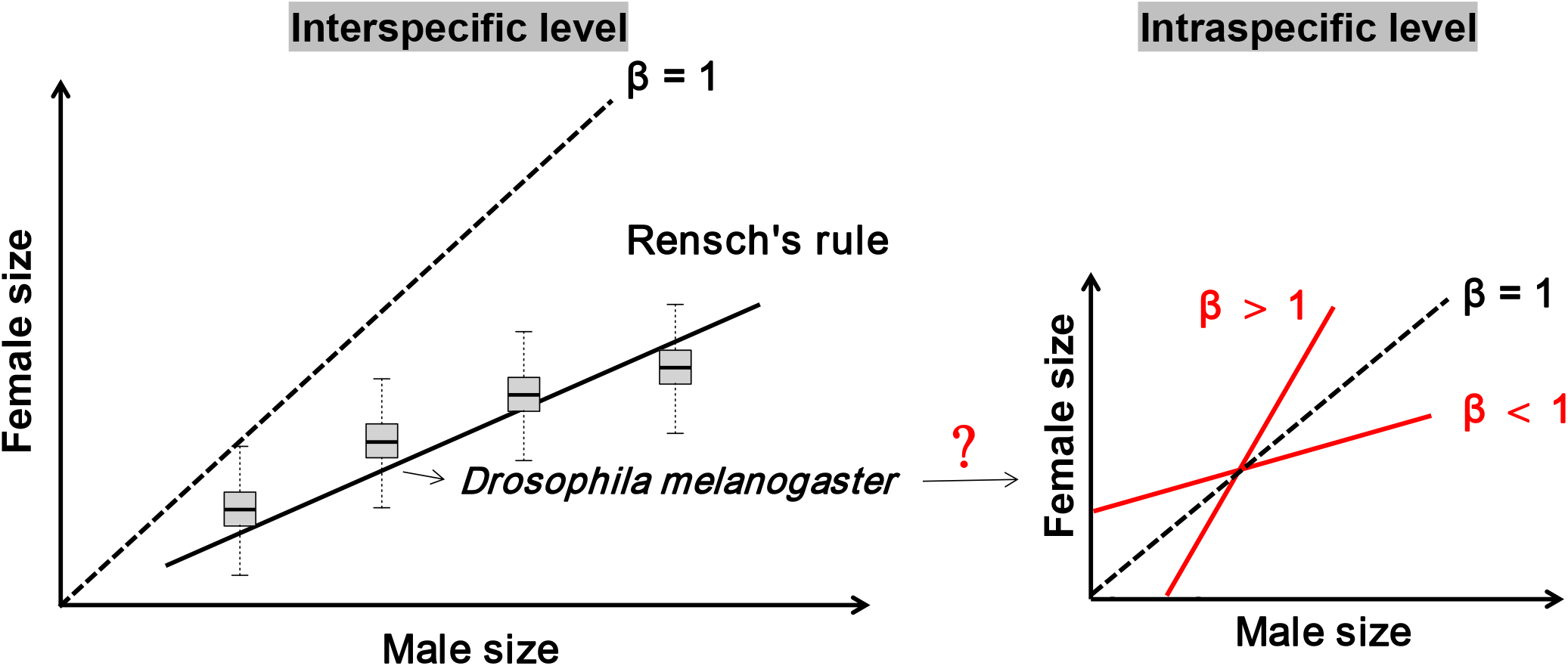
Conceptual illustration of potential scaling patterns of sexual size dimorphism (SSD). The dashed line (β = 1) represents the isometric expectation, where male and female body sizes increase proportionally. According to Rensch’s rule, SSD scales positively with body size when males are the larger sex, but scales negatively when females are larger (Left panel). At the intraspecific level, SSD may exhibit a reversed relationship (β > 1) or consistent trend (β < 1; Right panel). These variations in scaling patterns may arise from genetic differences, environmental factors, or ecological contexts within species.

In the present study, we investigated the intraspecific relationship between SSD and body size across inbred lines of the *Drosophila melanogaster* Genetic Reference Panel (DGRP). These lines provide standardized genetic backgrounds and an excellent system for testing how environmental conditions shape body size and SSD. Specifically, we aimed to: (1) test whether SSD in *D. melanogaster* follows Rensch’s rule at the intraspecific level; (2) determine whether sex-specific differences in growth rate contribute to intraspecific level divergence in SSD; and (3) evaluate whether temperature shifts intraspecific SSD by differentially altering male and female growth in *D. melanogaster*.

## MATERIALS AND METHODS

### *Drosophila* stocks and fly culture

We used the inbred lines of the *Drosophila melanogaster* Genetic Reference Panel (DGRP) to investigate the sexual dimorphism in size, development time and growth rate within species. The DGRP lines used in this study were originally derived from inseminated females collected in Raleigh, NC USA, and inbred by more than 20 generations of full sib mating to ensure genetic uniformity (Mackay et al., 2012). Prior to the experiments, flies were reared in standard laboratory conditions at 25 °C, under a 16:8 light/dark cycle at 70% relative humidity. The flies were cultured by food medium composed of 40g of dry yeast, 70g of cornmeal, 70g of sugar, 10g of agar, and 5 ml of propionic acid per liter of water to prevent contamination. All experiments were conducted at 25 °C and 28 °C chamber to examine the effects of temperature on sexual size dimorphism, development time and growth rate.

### Body size and development time measurements

In our study, *Drosophila melanogaster* reaches its peak egg-laying activity around 5 days post-eclosion. Accordingly, three pairs of 5-day-old virgin males and females from each DGRP line were introduced into individual vials (20 × 75 mm) containing 5 mL of standard medium. A minimum of ten replicates per line were initiated to ensure a sufficient sample size for subsequent measurement and statistical analysis. The flies were allowed to oviposit for 1 day, after which they were removed.

To assess development time, we recorded the emergence of each individual fly hourly, noting the time from egg laying to larval development. Newly emerged females and males were immediately anesthetized by freezing and weighed using a precision electronic balance (a precision of 0.000001g). The development time was determined by the larval developmental period as suggested by Teder (2014), while growth rate was calculated by dividing body mass of newly emerged adults by the duration of the larval stage.

### Statistical analysis

We quantified SSD of each DGRP line cultured at 25 °C and 28 °C as: SSD = (mean female size/mean male size) – 1 (Lovich & Gibbons, 1992). To examine sexual differences in development time and growth rate, we applied the same methodology. We refer to these differences as “dimorphism” in the context of this study. All dimorphism variables were and standardized (mean = 0 and SD = 1) to facilitate comparisons.

We first applied linear mixed models (LMMs) to evaluate sex-specific differences in these traits across 17 genotypes (11 genotypes at 28 °C). Body weight, development time, and growth rate were used as dependent variables. Sex and temperature were specified as fixed effects, whereas genotype was included as a random effect. The significance of random effects was assessed using likelihood ratio tests (LRTs), and model fit was compared based on Akaike Information Criterion (AIC) values, with lower AIC indicating a better-fitting model. All statistical analyses were performed using the *lme4* package (version 1.1-38) in R (Bates et al., 2015).

Because body size of both female and male individuals is estimated with error, we then employed the Standardized Major Axis (SMA) method with the *smatr* package (version 3.4.8) (Warton et al., 2012) to explore the allometric relationship between female and male body size within species. This allowed us to model the body size scaling relationship between the sexes across all genotypes. Additionally, we performed ordinary least squares (OLS) to assess the relationship between SSD and both sexual development time dimorphism (SDTD) and sexual growth rate dimorphism (SGRD). All statistical analyses were performed in R 4.4.1 (R Core Team, 2025).

## RESULTS

The linear mixed models (LMMs) revealed pronounced genotypic variation across all measured traits. Body weight, growth rate, and larval development time differed significantly among genotypes (see variance and AIC values in Table 1; *P* < 0.001 for all; Table 1). Strong sex effects were detected for body weight and growth rate, with females exhibiting faster growth and higher body weight than males (see estimates and F values in Table 1; *P* < 0.001 for both; Table 1), whereas larval development time did not vary between sexes (*P* = 0.38; Table 1).

Temperature exerted trait-specific effects. Body weight decreased markedly at 28 °C relative to 25 °C (*P* < 0.001; Table 1), and this reduction was sex-dependent (temperature × sex interaction: *P* < 0.001; Table 1), with females showing a larger decline than males (Figure 2a). Larval development time shortened at higher temperature (*P* < 0.001; Table 1), and the non-significant temperature × sex interaction (*P* = 0.71; Table 1) indicated that the acceleration in development was similar between the sexes (Figure 2b). In contrast, growth rate exhibited no main effect of temperature (*P* = 0.41; Table 1). However, a significant temperature × sex interaction (*P* < 0.001; Table 1) revealed opposite thermal responses between the sexes: growth rate decreased in females but increased in males from 25 °C to 28 °C (Figure 2c).

**Figure 2.**
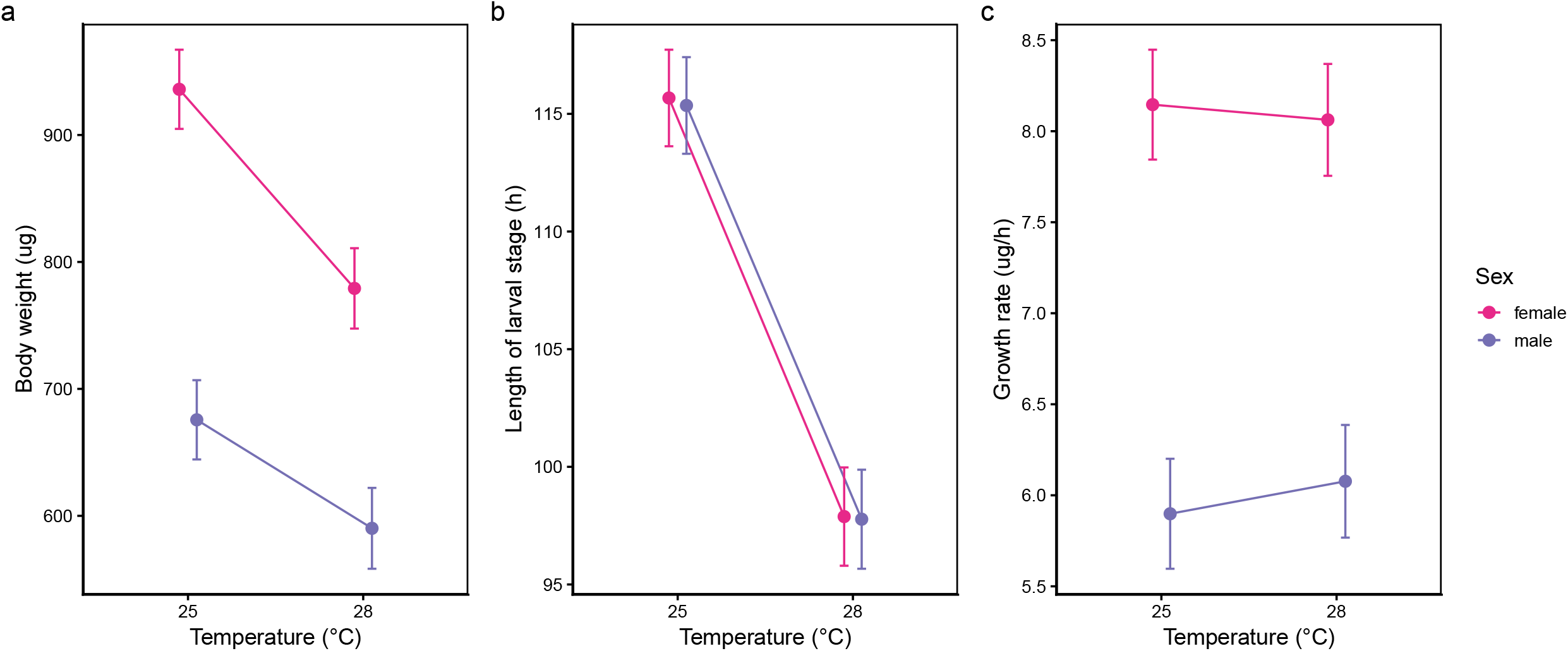
Effects of temperature (25 °C and 28 °C) and sex on (a) body weight, (b) length of larval stage, and (c) growth rate in *D. melanogaster*. Female measurements are shown in pink and male measurements in purple. Error bars denote standard errors. The temperature responses differ between sexes, indicating sex-specific reaction norms across traits.

In *D. melanogaster*, sexual size dimorphism (SSD) was consistently female-biased at both temperatures. Across genotypes, SSD increased with mean body size (Figure 3a; β > 1, *P* < 0.05), indicating a within-species reversal of Rensch’s rule. Similarly, sexual growth rate dimorphism (SGRD) became more pronounced with increasing mean growth rate (Figure 3b; β > 1, *P* < 0.01). By contrast, development time showed no significant sex differences at either temperature (*P* > 0.05). Elevated temperature reduced the magnitude of both SSD and SGRD (Figure 3). Moreover, SSD was strongly correlated with SGRD at both temperatures (Figure 4; 25 °C: estimate = 0.92 ± 0.10 SE; *R*^2^ = 0.83, *P* < 0.001; 28 °C: estimate = 0.90 ± 0.15 SE; *R*^2^ = 0.78, *P* < 0.001).

**Figure 3.**
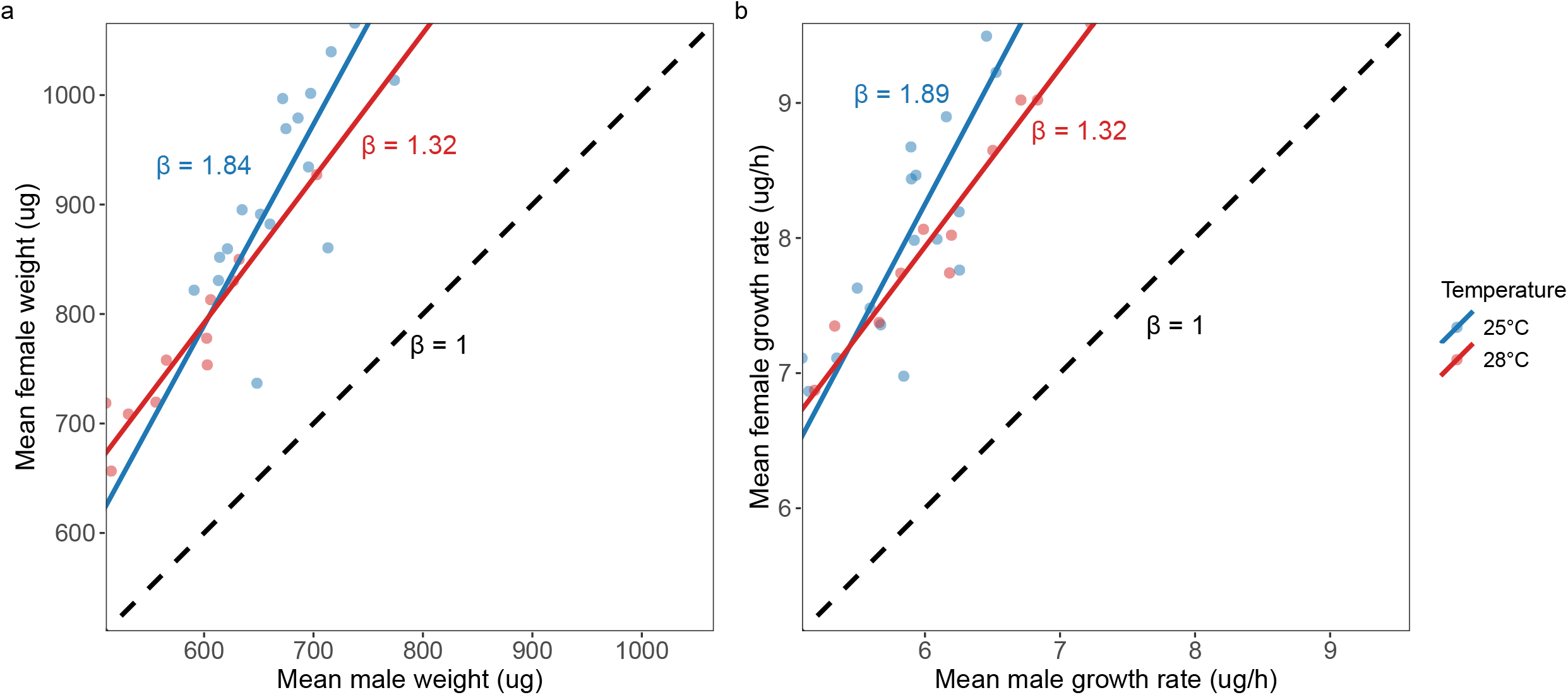
Allometric relationships between males and females for (a) body weight, and (b) growth rate in *D. melanogaster* reared at 25 °C and 28 °C. Standardized major axis (SMA) regression slopes (β) are provided; β = 1 indicates isometry, whereas β > 1 and β < 1 correspond to female-biased and male-biased allometry, respectively. The drakblue line represents the regression fit for flies reared at 25 °C, while the drakred line represents the regression fit for flies reared at 28 °C.

**Figure 4.**
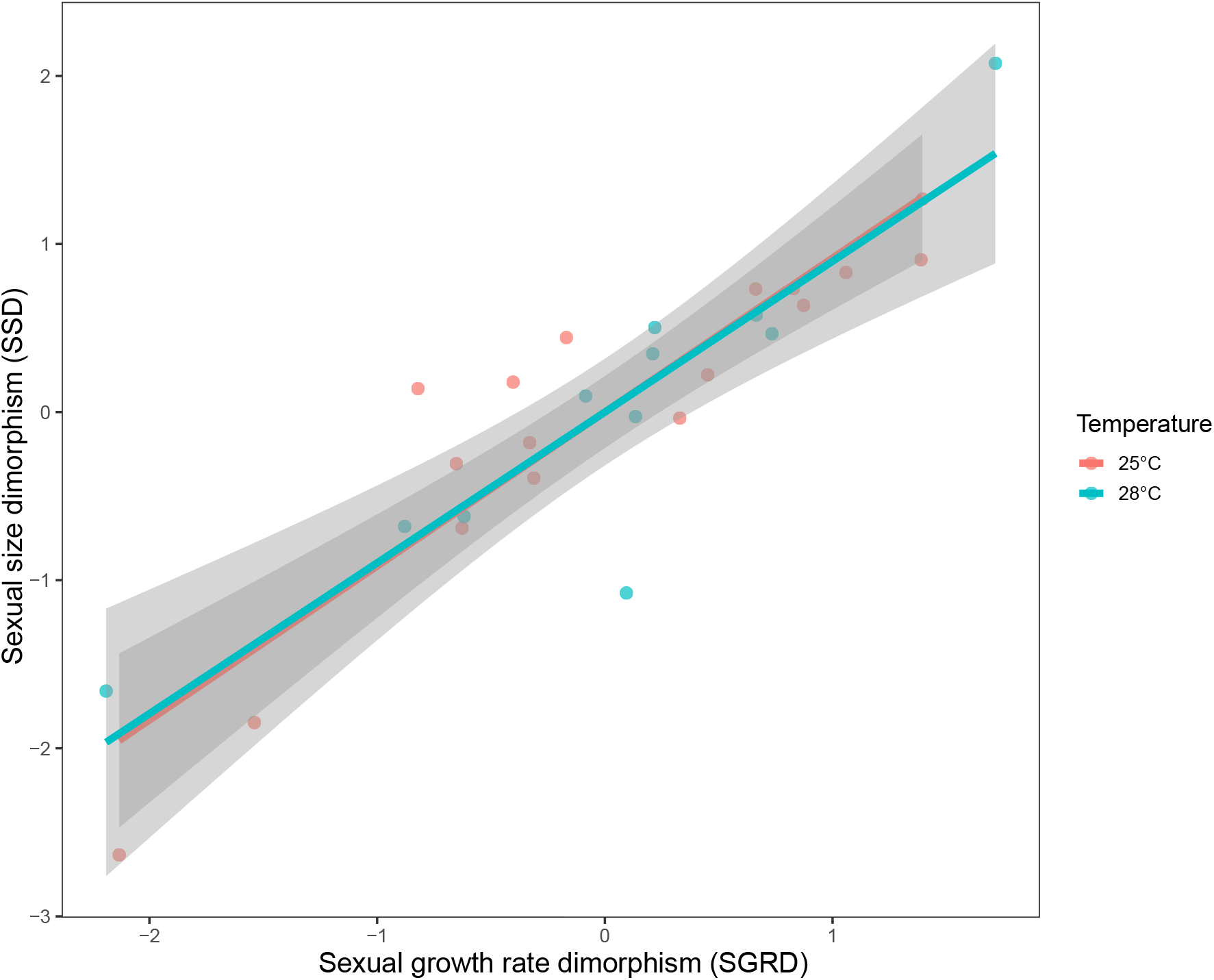
Associations between sexual size dimorphism (SSD) and dimorphism in growth rate (SGRD) in *D. melanogaster* reared at 25 °C and 28 °C. Each point represents a single isogenic line, with red points corresponding to flies reared at 25°C and green points corresponding to flies reared at 28 °C. Solid red and green lines represent significant ordinary least squares (OLS) regression fits for 25 °C and 28 °C treatments, respectively. The shaded gray area represents the 95% confidence interval for each regression fit.

## DISCUSSION

In this study, we explored the intraspecific scaling relationship between sexual size dimorphism (SSD) and body size in *D. melanogaster* using inbred *Drosophila melanogaster* Genetic Reference Panel (DGRP) lines under two thermal conditions. Our study yielded three major findings. First, females consistently outgrew males, resulting in a pronounced female-biased SSD. Second, SSD increased with overall body size, representing a reversal of Rensch’s rule at the intraspecific level. Third, the variation in SSD was primarily driven by faster growth of female individuals rather than differences in development time between male and female individuals.

Taken together, we found that SSD in *D. melanogaster* increases with body size despite the persistent female-biased dimorphism. Our results provide compelling evidence for a reversal of Rensch’s rule, i.e., SSD decrease with body size when females are bigger than males (Abouheif & Fairbairn, 1997; Fairbairn, 1997). Interestingly, previous investigations at the interspecific level of drosophilid species found the SSD and average body size conform to Rensch’s rule (Teder & Tammaru, 2005; Blanckenhorn et al., 2007). This scale-dependent discrepancy suggests that the selective forces shaping SSD evolution may shift or interact differently across evolutionary levels. Sex-specific selection, especially fecundity selection on females, can promote disproportionate increases in female size relative to males, thereby amplifying female-biased SSD at intraspecific level (Shine, 1988; Honek, 1993). In contrast, across species, shared genetic and developmental architecture can couple male and female size evolution, limiting their independent divergence and generating interspecific SSD-body size relationship described by Rensch’s rule (Abouheif & Fairbairn, 1997; Badyaev, 2002). Overall, these findings highlight the need to integrate both intraspecific and interspecific perspectives to fully understand the evolutionary of SSD and the applicability of Rensch’s rule.

Furthermore, we provide evidence that sex-specific differences in growth rate contribute to the proximate basis of female-biased SSD in *D. melanogaster*. Although males and females exhibited similar developmental durations, female larvae accumulated biomass more efficiently, resulting in larger adult body sizes. Thus, female biased SSD of *D. melanogaster* is driven primarily by growth rate rather than prolonged development. This is consistent with observations in other arthropods, regardless of conformity to Rensch’s rule, where SSD has often been attributed to sex-specific growth rate differences (Blanckenhorn et al., 2007; Stillwell & Davidowitz, 2010). Body size is commonly achieved at the cost of prolonged development under constrained growth conditions (Roff, 1992). Higher growth rate could allow females to realize fecundity gains from larger size while limiting the survival costs of extended juvenile development, including increased exposure to natural enemies (Gotthard et al., 1994; Davidowitz & Nijhout, 2004). Furthermore, our thermal treatment experiments further revealed that growth rate is sensitive to environmental change and can modulate the degree of SSD. At elevated temperatures, female growth rate declined, whereas male growth rate increased, leading to a reduction in SSD. These results suggest that males and females exhibit distinct reaction norms, resulting in asymmetric plasticity to temperature (Teder et al., 2022). Climate change may therefore impose sex-specific constraints on growth, and affect the adaptive potential of natural populations.

Although males and females exhibited different growth responses across temperature treatments, both sexes showed a similar larval development time across both temperatures. This convergence may reflect selection to limit sex-specific differences early in development, reducing the chances of mismatches in adult emergence and mating availability (Thompson et al., 2023). In species like *D. melanogaster*, where the adult lifespan is short and resources are available for only a brief period, even slight shifts in emergence timing can impose significant fitness costs that females may have fewer mating opportunities, while males may miss access to receptive females (Morbey & Ydenberg, 2001). Maintaining closely aligned developmental schedules thus helps preserve effective mating synchrony, a factor especially critical under intense scramble competition in which delayed emergence substantially diminishes male reproductive success (Davies & Saccheri, 2015; Thompson et al., 2023). Consequently, although growth rates display sex-specific plasticity, the conservation of similar development time likely represents an adaptive strategy shaped by sexual selection to stabilize mating interactions and sustain reproductive output across variable thermal environments.

Finally, while our study provides novel insights into reversed Rensch’s rule at the intraspecific level, several limitations should be acknowledged. First, we examined only two thermal regimes, assessing a broader gradient of temperatures would allow a more comprehensive understanding of the plasticity of SSD. Second, our experiments were conducted under controlled laboratory conditions with standardized resources and minimal competition, which may not fully capture the ecological complexity of natural environments. Therefore, future research should investigate SSD across a wider range of environmental contexts, including resource limitation and interspecific interactions, to better reflect the ecological and evolutionary forces shaping sexual dimorphism. Finally, integrating genomic or transcriptomic analyses of the DGRP lines could provide valuable insights into the genetic underpinnings of sex-specific growth plasticity and its contribution to the evolution of SSD.

## Figure legends

**Table 1.** The results of linear mixed models (LMMs) examining the effects of sex and temperature on body weight, larval development time, and growth rate among *D. melanogaster* genotypes.

